# Automatic subtyping of individuals with Primary Progressive Aphasia

**DOI:** 10.1101/2020.04.04.025593

**Authors:** Charalambos Themistocleous, Bronte Ficek, Kimberly Webster, Dirk-Bart den Ouden, Argye E. Hillis, Kyrana Tsapkini

**Affiliations:** Johns Hopkins School of Medicine, Baltimore, MD 21287, USA; Arnold School of Public Health, University of South Carolina, Columbia, SC, USA

**Keywords:** Primary Progressive Aphasia, Classification, Natural Language Processing, Machine Learning

## Abstract

**Background:** The classification of patients with Primary Progressive Aphasia (PPA) into variants is time-consuming, costly, and requires combined expertise by clinical neurologists, neuropsychologists, speech pathologists, and radiologists.

**Objective:** The aim of the present study is to determine whether acoustic and linguistic variables provide accurate classification of PPA patients into one of three variants: nonfluent PPA, semantic PPA, and logopenic PPA.

**Methods:** In this paper, we present a machine learning model based on Deep Neural Networks (DNN) for the subtyping of patients with PPA into three main variants, using combined acoustic and linguistic information elicited automatically via acoustic and linguistic analysis. The performance of the DNN was compared to the classification accuracy of Random Forests, Support Vector Machines, and Decision Trees, as well as expert clinicians’ classifications.

**Results:** The DNN model outperformed the other machine learning models with 80% classification accuracy, providing reliable subtyping of patients with PPA into variants and it even outperformed auditory classification of patients into variants by clinicians.

**Conclusions:** We show that the combined speech and language markers from connected speech productions provide information about symptoms and variant subtyping in PPA. The end-to-end automated machine learning approach we present can enable clinicians and researchers to provide an easy, quick and inexpensive classification of patients with PPA.

## Introduction

Primary Progressive Aphasia (PPA) is a progressive neurological condition that is characterized by a gradual deterioration of speech and language. The term PPA was coined by Marsel Mesulam in his seminal paper in 1982 [1], almost a century after Arnold Pick’s early observations in 1892 of progressive and irreversible forms of dementia affecting language [2]. In patients with PPA, there is substantial symptom variability as a result of neurodegeneration affecting different brain areas. In an organized attempt to categorize speech and language deficits in patients with PPA, a team of experts put forth specific guidelines in a consensus paper confirming the variability of the syndrome and providing criteria for classification of patients with PPA into three main variants: nonfluent variant PPA (nfvPPA), semantic variant PPA (svPPA), and logopenic variant PPA (lvPPA) [3].

Patients with nfvPPA are characterized by effortful speech with sound errors and distortions and impaired production and perception of syntactically complex sentences. These patients have predominant damage in the left inferior frontal lobe, which is usually associated with frontotemporal dementia pathology and tauopathies. Patients with svPPA are characterized by difficulties in confrontation naming, single-word comprehension, and impaired semantic memory of familiar objects that often results in ‘empty speech’ (i.e., output without meaning).

Their predominant damage is in the left and right anterior temporal lobes. Patients with svPPA are associated with having TAR DNA-binding protein 43 (TDP-43) pathology. Patients with lvPPA are characterized by difficulties in word retrieval, repetition of long words and phrases, and phonological errors. Their predominant damage is in the left posterior temporal and inferior parietal areas, left anterior hippocampus and precuneus, and their underlying pathology is usually linked to Alzheimer’s Disease (AD).

Since patients with the same PPA variant share common linguistic deficits, PPA variants can inform the type of language therapy provided, such as targeting word retrieval, sentence formulation strategies, and addressing oral apraxia. However, the task of subtyping individuals with PPA into variants is time consuming, arduous, and requires combining evaluations by clinical neurologists, neuropsychologists, speech pathologists, and radiologists. As insurance companies typically cover only limited therapy sessions, and the condition is progressive, the decision about the type of variant must be made quickly. Therefore, there is a dire need for an easy, quick, and accurate evaluation, consistent with the established criteria and sensitive to the speech and language deficits associated with each variant. An automatic evaluation system based on machine learning has the potential to save clinicians time and provide important information with respect to PPA variant and clinical treatment.

This study shows a machine learning model based on Deep Artificial Neural Networks (DNNs) that can subtype patients with PPA into variants with high classification accuracy. This automated approach offers diagnosis tailored to specific individuals, using information from connected speech productions elicited from a picture description task—a naturalistic task that takes fewer than two minutes to administer. This model has two main advantages. First, information about the PPA variant of a patient can enable clinicians to make better decisions about tests and therapy batteries that can elicit optimal therapy results for individuals with PPA. Thus, knowledge about the PPA variant will enable assessment and treatment that is tailored to the specific individual with PPA [4]. Second, by knowing the PPA variant, clinicians can share knowledge about successful treatment solutions that target specific subpopulations of individuals with PPA, which can be comparable across individuals and clinics. Treatments that lead to improved speech and language performance in individuals with one variant of PPA can be enhanced and applied to new individuals identified with the same variant, and unsuccessful treatments can be disregarded.

The machine learning model employs information from connected speech productions. Connected speech can convey a striking amount of information about PPA variants and is readily available through a simple picture description task [5]. For instance, it can inform patterns of articulation (e.g., vowel production), prosody and grammar (e.g., morphology and syntax), and lexical retrieval. Earlier studies showed that the relative vowel duration from a polysyllabic word repetition task can distinguish patients with nfvPPA and lvPPA [6]. Independent evidence on the role of vowels in PPA-variant classification comes from studies of paraphasias [7] and from cross-sectional studies accompanied by lesion-symptom mapping [8]. Patients with svPPA use more pronouns than nouns [9]. Patients with nfvPPA are characterized by impaired production of grammatical words, such as articles, pronouns, etc., but have fewer difficulties using words with lexical content than individuals with svPPA [3]. The use of linguistic features has also been shown to distinguish PPA from patients with AD. For example, a current study using morphosyntactic and semantic features in combination with other features, such as the age of acquisition of words, and age of acquisition of nouns and verbs, resulted in a good classification accurac of nfvPPA from AD and healthy individuals [10]. Combined morphosyntactic features can provide additional information about patterns of speech and language productions in individuals with PPA. For example, the *noun-verb ratio* can reveal whether an individual with PPA shows preference towards nouns or verbs, which may help to subtype nfvPPA from patients with svPPA [9, 10]; similarly, as patients with svPPA are impaired in noun naming, the *noun-pronoun ratio* can distinguish patients with svPPA from patients with another variant [9]. Adding such information about morphosyntax can enable the machine learning model to make better predictions.

Current neural networks enable the representation and modeling of complex problems, including natural language processing, speech recognition, image recognition, and machine translation [11]. They are the result of the rapid advancements that have taken place over the last few years. What makes them a model of choice for this paper is a number of recent developments in their architecture that enable exceptional learning of abstract representations. The early neural network developed by McCulloch and Pitts in 1943 could compute logical operations but could not learn [12]; learning was made possible almost a decade later, by networks proposed by Hebb [13] and Rosenblatt [14] and through the formulation of backpropagation, which enabled learning in multi-layered networks [15]. Current state-of-the art neural networks not only learn but outperform other methods, including Support Vector Machines (SVMs), Random Forests (RF), and Decision Trees (DTs) [16, 17] in most classification problems. Neural networks are not homogeneous, as different architectures exist, such as feed-forward networks, recurrent convolutional networks, and others [e.g., 11, 17, 18, 19, 20]. This allows for flexibility in addressing different learning problems in different learning scenarios, such as classification, regression, or sequence modeling, in supervised, unsupervised, or reinforcement learning. Given the successes of DNNs, we can expect that they can learn multiple levels of representation required to classify the different variants of PPA, identify patterns from the acoustic and linguistic structure, and enable the development of automatic systems for PPA subtyping, thus resulting in improved targeting of treatment.

The aim of this study is twofold: (1) to provide a machine learning model that can automatically subtype PPA and offer diagnosis tailored to specific individuals, using information from connected speech productions elicited by a simple picture description task that takes fewer than two minutes to administer, and (2) to contribute to our current understanding of PPA variants and their differences in speech and language characteristics.

We employed data from connected speech productions via a simple and widely used picture description task: the Cookie Theft description from the Boston Diagnostic Aphasia Examination [21]. The task was administered to participants with PPA as part of a large clinical trial (NCT:02606422) during baseline evaluation sessions. Picture description productions were automatically transcribed and segmented into words, vowels, and consonants. Acoustic features were then extracted from vowel productions. We have excluded consonants, as they require a different type of feature analysis depending on their manner of articulation (stops, fricatives, sonorants, etc.) and may have an additional computational cost (e.g., dimensionality, longer processing and extraction). The whole text from transcribed picture descriptions of participants with PPA was analyzed using automatic morphsyntactic analysis that provides the part of speech (POS) for each word in the text. Acoustic and linguistic measurements were combined to provide the predictors for the machine learning analysis. To achieve the subtyping of PPA variants, we developed DNN architectures that were trained on the combined set of acoustic and linguistic measurements. Results for the model comparison of DNNs with RF, SVMs, and DTs are also reported. The methodology part details the complex procedure of designing and evaluating an end-to-end machine learning model based on neural networks. However, we need to stress out that this process is conducted only once. The potential users of the model, speech/language therapists, clinicians, etc. will not need to train or evaluate the machine learning model again but they will upload the recording with the cookie-theft into the system and the model will provide the PPA variant automatically.

## Methods

### Data Collection

Trained clinicians or clinician assistants recorded 44 individuals with PPA during baseline evaluation sessions (**Table 1**). All participants had a diagnosis of PPA from an experienced neurologist, a history of at least two years of progressive language deficits with no other etiology (e.g., stroke, tumors, etc.), and relatively preserved memory as shown from the general Clinical Dementia Rating (CDR) Scale [22]. Participants were also right-handed and native speakers of English. Differential diagnosis of individuals with PPA and PPA variant subtyping was based on Magnetic Resonance Imaging (MRI) results, clinical and neuropsychological examination, and speech and language evaluations following the consensus criteria by Gorno-Tempini, et al. [3] by experienced neurologists. Specifically, 9 participants were subtyped as svPPA, 16 as lvPPA, and 19 as nfvPPA.

**Table 1.** Demographic information of the participants for each PPA variant (for age, education, onset of the condition in years, language severity, and total severity, the mean and the standard deviation in parenthesis is provided; language severity and total severity correspond to the Behaviour-Comportment-Personality and Language domains of the FTLD-CDR [22].

| Variant | svPPA | lvPPA | nvPPA |
| --- | --- | --- | --- |
| Female | 5 | 8 | 7 |
| Male | 4 | 8 | 12 |
| Total Patients | 9 | 16 | 19 |
| Age | 66.59 (6.06) | 67.93 (7.55) | 69.07 (5.57) |
| Education | 16.30 (1.92) | 16.92 (2.24) | 16.42 (1.37) |
| Onset years | 6.48 (2.31) | 3.88 (3.23) | 3.49 (1.80) |
| Language severity | 2.27 (0.56) | 1.39 (0.75) | 1.77 (0.48) |
| Total severity | 7.75 (4.36) | 4.98 (2.82) | 6.04 (3.10) |

Table 1 provides biographical demographic information of the participants for each PPA variant. One-way ANOVA tests showed that there were no significant differences between variants for gender (*F*=5.82, *df*=2, *p*= 0.09), age (*F*=0.354, *df*=2, *p*=0.705), education (*F*=0.162, *df*=2, *p*=0.853), language severity (*F*=0.154, *df*=2, *p*=0.86) and total severity (*F*=1.162, *df*=2, *p*=0.33). We used the revised Frontotemporal Dementia Clinical Dementia Rating (FTD-CDR) Scale to rate severity of PPA [22]. Severity was calculated by three independent raters who scored each item for each participant based on face-to-face interaction with the participant and family, along with language and cognitive evaluations. Total severity and language severity scores were the result of the consensus between the three raters. Data collection was conducted as part of a clinical trial on Transcranial Direct Current Stimulation for Primary Progressive Aphasia at Johns Hopkins University (NCT:02606422). All participants provided informed consent.

### Data preprocessing

Recordings from the Cookie Theft picture description task were saved in mono waveform audio file format (wav files) at a 16000 Hz sampling frequency. The following three preprocessing pipelines were developed to analyze the acoustic and linguistic properties and generate the classification data.

#### Pipeline 1 Audio transcription and segmentation

The sounds were processed using Themis, a python program developed in-house that provides a text file with the audio transcription of each word and segment—vowel, consonant, pause—and a table that contains the times (onset time and offset time) of each word and segment [9, 23, 24]. The table was converted into Praat TextGrid files for processing in Praat automatically [25] pause duration was calculated during segmentation from the automatic alignment system.

#### Pipeline 2 Audio processing

A second pipeline in Praat software for speech analysis enabled the extraction of acoustic information for the segmented vowels. Specifically, the following acoustic properties were measured:

i. *Vowel formants*. Formant frequencies from first formant frequency to the fifth formant frequency were measured at the 25%, 50%, and 75% mark of vowel duration.
ii. *Vowel duration*. Vowel duration was measured from the onset to the offset of the first and second formant frequencies. Other acoustic cues, such as the rise of the intensity, were employed to determine the left and right vowel boundaries.
iii. *Fundamental frequency*. (*F*0). We calculated the mean *F*0, minimum *F*0, and maximum *F*0 for each vowel production. *F*0 calculation was conducted using the autocorrelation method.
iv. *H1–H2, H1–A1, H1–A2, H1–A3*. Harmonic and spectral amplitude measures were extracted from the vowels using Praat.

Overall, we employed the following 40 predictors: vowel duration, pause duration, and the first five formant frequencies measured at three locations inside the vowel at the 25%, 50%, and 75% of the vowel duration, voice quality features (H1-H2, H1-A1, H1-A2, H1-A3), measures of *F*0 (Minimum *F*0, Mean *F*0, Maximum *F*0), and POS ratios and means (noun/verb ratio, noun/adjective ratio, noun/adverb ratio, noun/pronoun ratio, verb/adjective ratio, verb/adverb ratio, verb/pronoun ratio, adjective/adverb ratio, adjective/pronoun ratio, adverb/pronoun ratio, mean noun, mean pronoun, mean verbs, mean adjective, mean adverb).

#### Pipeline 3 Morphosyntactic analysis

A third pipeline processed transcripts morphosyntactically. It conducted an automatic morphosyntactic analysis using the TextBlob python library [26]. Measurements of characters, words, characters per word, etc., were calculated from the tokenized and parsed output, and the ratio of each part of speech per total number of words and the ratio between two part of speech categories, i.e., the noun-verb ratio, noun-adjective ratio, noun-adverb ratio, noun-pronoun ratio, verb-adjective ratio, verb-adverb ratio, verb-pronoun ratio, adjective-adverb ratio, adjective-pronoun ratio, and adverb-pronoun ratio were calculated.

The outputs of the three pipelines were combined into a single comma-separated values (CSV) file that was employed as an input for the machine learning models.

### Neural Networks

Appendix 1 explains the design of the Neural Network and provides the necessary details for replicating its design and evaluating its architecture, allowing our approach to be compared to other modeling work. Providing the appropriate details on the architecture and implementation of neural networks necessitates some use of jargon, which we attempted to limit only to the Appendix. We also cite the necessary related works that clarify this technical jargon. In the following we discuss the model comparison process and the evaluation.

#### Comparing the performance of Neural Networks to other machine learning models

To estimate the performance of the DNN, we show how three machine learning models are compared to the DNN model. We employed support vector machines (SVM) [27], random forests (RF) [28, 29], and decision trees (DT) as these are often used in medical studies [30].

i. DTs provide a multiclass classification of patients with three PPA variants by splitting the data using the measurement that best explains the variability of the data. For example, if the ratio of nouns explains most of the variation, the decision tree will split the data in two branches based on the ratio of nouns and will repeat the process multiple times exhaustingly, i.e., up to the point there are no data. One major advantage of DTs is that trees can be visualized and can provide an understanding of the structure of the data and the exact decisions that the model made for the classification. Nevertheless, DTs are often prone to overfitting, as they create long and complicated trees that may not generalize very well to unknown data. Even though there are methods to control overfitting, such as by *pruning* the lower branches of the tree that explain very little of the remaining variation, these methods are not always optimal.
ii. SVMs classify multidimensional data, using a separating hyperplane that organizes the data points into classes. The data points that delimit the hyperplane are called *support vectors* and the separating hyperplane is considered a classification *machine*. One advantage of SVMs is that they can provide good classification results. One disadvantage of SVMs is that the optimization of their tuning parameters (a.k.a., hyperparameters) can be complex and time consuming.
iii. RFs are similar to DTs, but, unlike DTs, they are ensemble models, i.e., they fit several decision trees on the measurements collected and combine them using an ensemble measures such as the mean, to improve the accuracy of the model. RFs can address the overfitting that often takes place in the case of DTs.

### Comparison to Human Raters

Three trained speech-language pathologists, who were not involved in data collection, provided a classification by listening to Cookie Theft recordings from nine participants who were employed for the training of this model (three participants from each variant). Their responses were evaluated using the information that was provided by the combined neuro-psychological examination, MRI, etc.

## RESULTS

The Cookie Theft picture description recordings were analyzed to elicit measures of speech and language from patients with PPA. These measures were then employed to train a DNN, along with three other machine learning models, namely a Random Forest, a Support Vector Machine, and a Decision Tree, to provide comparative results for estimating the performance of the DNN. All machine learning models were trained and evaluated using an 8-fold cross-validation method. “True class” variant diagnoses were given by experienced neurologists. Table 2 shows the results from the 8-fold cross-validation method. Overall, the DNN provided 80% classification accuracy and outperformed the other three machine learning methods. The Support Vector Machines had the worst performance in the cross-validation task with 45% classification accuracy (see in Table 4 panel a). Random Forests (RFs) provided a 58% classification accuracy (see in Table 4 panel b), followed by the Decision Tree model (DT) with 57% classification accuracy (see in Table 4 panel c).

**Table 2.**
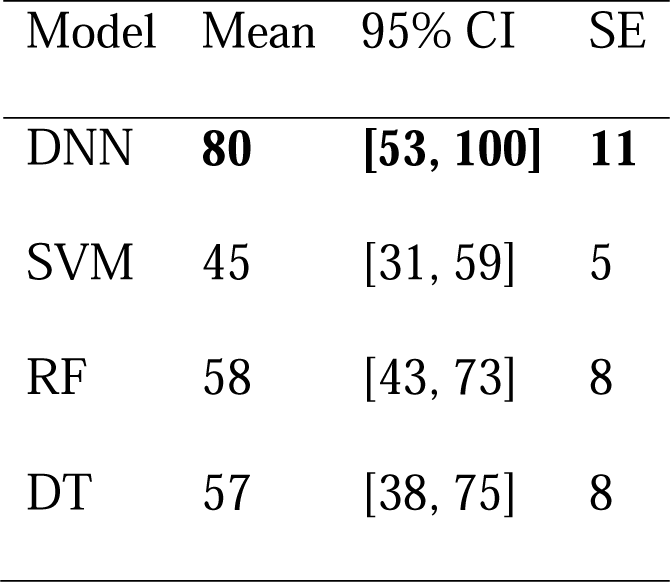
Results from eight-fold cross-validation for the Deep Neural Network (DNN), Support Vector Machines (SVM), Random Forest (RF), and Decision Tree (DT). Shown is the mean cross-validation accuracy, the 95% Confidence Intervals (95% CI) and the standard error (SE).

| Model | Mean | 95% CI | SE |
| --- | --- | --- | --- |
| DNN | **80** | **[53, 100]** | **11** |
| SVM | 45 | [31, 59] | 5 |
| RF | 58 | [43, 73] | 8 |
| DT | 57 | [38, 75] | 8 |

The confusion matrix shown in Table 3 and 4 was calculated by summing the 8 confusion matrices produced during cross-validation for the DNN. The DNN provided improved identification of patients with lvPPA and nfvPPA with respect to svPPA. Patients with lvPPA were identified 95% correctly; 5% of patients with lvPPA were identified as nfvPPA. Patients with svPPA were correctly identified in 65% of the cases; 30% of patients with svPPA were misclassified as lvPPA, and 6% as nfvPPA; 90% of patients with nfvPPA were correctly identified and 10% were classified as svPPA.

**Table 3.** Normalized confusion matrix from the evaluation of the Deep Neural Network. The confusion matrix provides the sum of scores from the 8-fold cross-validation test, showing the actual values vs. the predicted values.

|  |  | Predicted Class |  |  |
| --- | --- | --- | --- | --- |
|  |  | svPPA | lvPPA | nfvPPA |
| True Class | svPPA | 64 | 30 | 6 |
|  | lvPPA | - | 95 | 5 |
|  | nfvPPA | 10 | - | 90 |

**Table 4.**
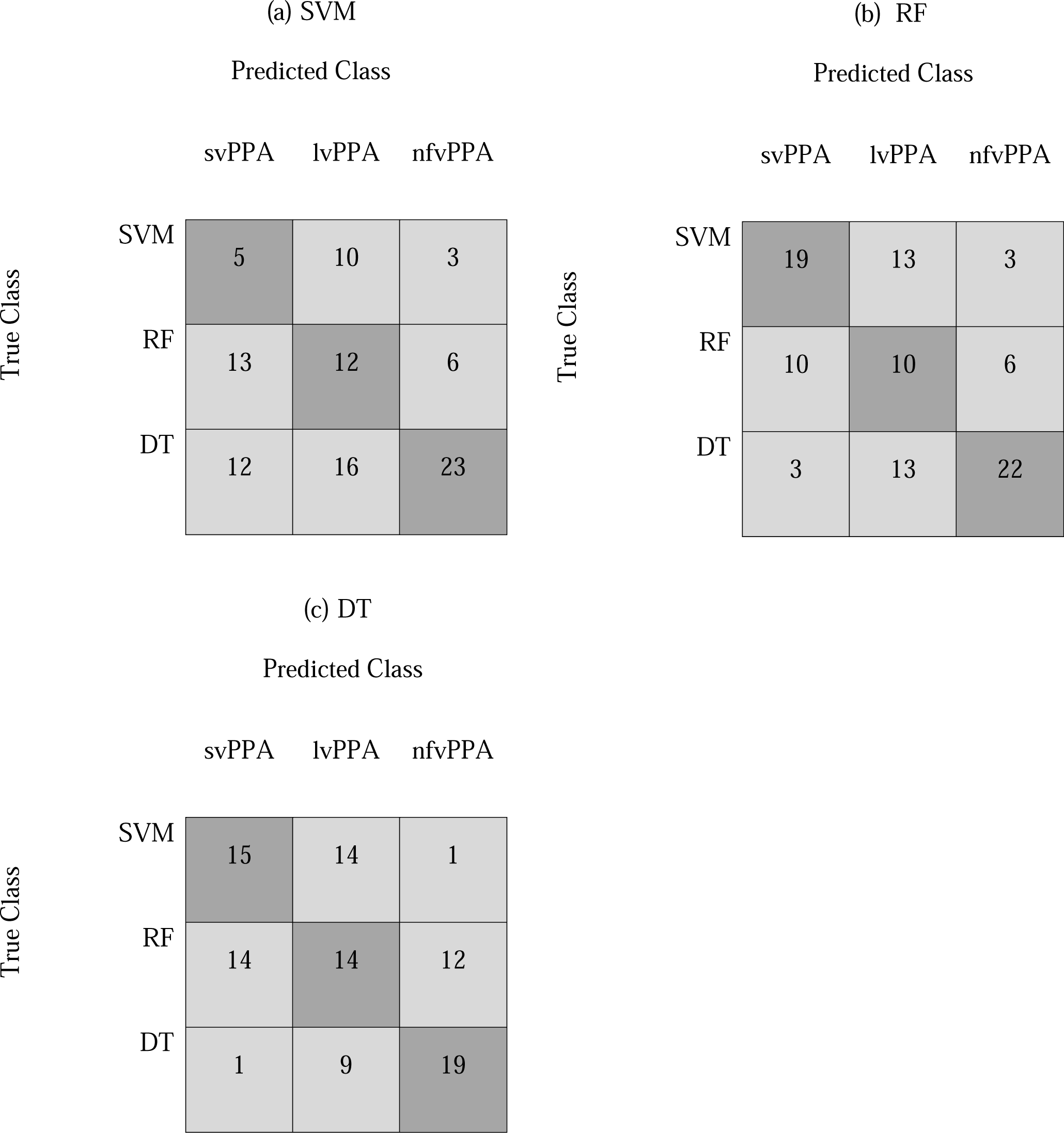
Normalized confusion matrix created by summing scores across cross-validation scores from the 8-fold cross-validation tests for SVM (Panel a), RF (Panel b), and DT (Panel c); matrices show the predicted vs. actual values from the evaluation.

To estimate the performance of the DNN, we also compared its accuracy with the classification performance of three trained speech-language pathologists (SLPs) who did not work with the patients who participated in the study. The SLPs’ classifications were based solely on the two-minute Cookie Theft samples. Their responses were compared to the gold standard combined subtyping that employs neuropsychological tests, imaging, language evaluation, etc. The three SLPs displayed significant variation in their classification scores of patients’ variants with mean classification accuracy 67% (SD=11). One SLP’s correct identification of the PPA variant of patients was just above average (5/9) 56%, followed by one who had (6/9) 66%, classification accuracy, and the highest classification accuracy reached (7/9) 77.77%. Overall, using the same evaluation data, the DNN provided more accurate results than SLPs and at a much faster rate.

## Discussion

Manual subtyping of patients with PPA is time-consuming and requires a high degree of expertise on classification criteria, costly scans, and lengthy evaluations. In this study, neural networks were trained on acoustic and linguistic predictors derived from descriptive-speech samples from patients with different PPA variants. The gold-standard classification of PPA patients was based on expert clinical and neuropsychological examinations, MRI imaging, and speech-language evaluations following established consensus criteria [3]. All models were trained 8 times in an eight-fold cross-validation model evaluation method. The output of the DNN was compared to the performance of three other machine learning models, namely Random Forests, Decision Trees, and Support Vector Machines, as well as with human auditory classification by experienced speech-language pathologists. The DNN achieved an 80% classification accuracy and outperformed the three other machine learning models. Acoustic and linguistic information from a short picture description task enables highly accurate classification of patients with PPA into variants.

Importantly, the model outperformed clinicians when provided the same information (only Cookie Theft picture descriptions). These results illustrate three important conclusions: (a) a minimal amount of acoustic and linguistic information from connected speech has great discriminatory ability, providing an identification fingerprint of patients with different PPA variants when used in a DNN model, (b) the DNN can simultaneously perform classification of all three PPA variants, and (c) the present automated end-to-end program may significantly help both the expert clinician by confirming the variant diagnosis, as well as the novice or less experienced clinician by guiding the variant diagnosis.

An unexpected finding was the improved classification results for the patients with lvPPA. The DNN model performed better than other machine learning models for lvPPA used by Hoffman, et al. [31] and Maruta, et al. [32], for example. Hoffman, et al. [31] employed unsupervised classification methods and analyzed results from linguistic (e.g., hesitations, phonological errors, picture-naming scores, single-word comprehension, category fluency scores, written competence) and non-linguistic (cube analysis, paired associate learning, etc.) neuropsychological evaluations. They found that participants with lvPPA were not identified as a separate group but were mixed with other participants in both linguistic and non-linguistic tasks [31]. Another study by Maruta, et al. [32] using a combination of measures from language and neurophysiological assessments in Portuguese discriminates individuals with svPPA from nfvPPA but not individuals with nfvPPA and svPPA from lvPPA [32].

The DNN machine learning model provides superior results compared to human auditory classification from three trained speech-language pathologists who were asked to provide a classification by listening to Cookie Theft productions. They listened to the same recordings that were used for training the network and scored lower than the DNN. Also, clinicians differed considerably in their judgments and often had to listen multiple times to the recordings to provide a judgment about the variant. The clinician with the highest classification accuracy reported that she had to listen several times for the patients who had “mild effects.” Overall, human PPA subtyping based on the same information provided to the machine learning model (Cookie Theft descriptions) can be a very difficult task for clinicians, including SLPs, because of different training and experience levels, as well as the use of different criteria for PPA subtyping. Moreover, speech samples are often hard to hear in PPA patients requiring frequent playback and attentive listening.

The limited amount of data employed constitutes the main limitation of our study. Although 44 patients with PPA is a very substantial number for a rare syndrome such as PPA, increasing the training data will enable the DNN to identify patterns between acoustic and linguistic predictors that characterize each variant with increased confidence. In fact, during the evaluation of machine learning models, it became evident that the amount of data in the training set has a significant impact on model accuracy. By increasing the overall data sample and obtaining data from more patients, the model’s accuracy will improve. A second limitation is inherent to the task used for eliciting connected speech samples, i.e., the Cookie Theft picture description. This task constrains speech production both acoustically and with respect to the required grammar. Patients provide primarily declarative intonational patterns, whereas questions, commands, etc. are not elicited. Also, picture-description tasks tend to elicit actions in the present tense, and sentences with factual content rather than wishes, commands, embedded sentences and other more complex structures. By contrast, other tasks, such as personal story telling or naturalistic conversation, have the potential to provide more informative speech and language output. Future classification work is likely to benefit from machine learning models trained on simultaneous classification of PPA variants using multifactorial predictors from a variety of discourse settings and conversations. Also, in our future research we plan to develop neural network models that distinguish patients with PPA from healthy controls (see for example [23]). A machine learning model trained on healthy controls that can distinguish patients with PPA from individuals with similar sociolinguistic characteristics (e.g., age, education, etc.) without PPA can complement the subtyping process.

## Acknowledgements

We thank our funding resources for their support: the Science of Learning Institute grant ‘Effects of tDCS in PPA’ from Johns Hopkins University to KT, and NIH/NIDCD R01 DC14475 to KT. We thank Ms. Olivia Hermann for her assistance in this work.

## Author contributions

C.T. conducted the acoustic analysis of the materials, designed the machine learning models, and wrote the first draft of the paper. K.T. led the trial where the data were acquired; B.F. and K.W. collected the acoustic materials. Subsequently, C.T., B.F., K.W., D.O., A.H., and K.T. revised the text. All authors approved the final version.

## Data and materials availability

The code is publicly available at https://github.com/themistocleous/ppa_classification; the data are not publicly available because they can potentially be employed to identify the participants.

## Appendix 1.

### Neural Network Architecture

The Appendix provides a description of the architecture of the Neural Network and the design of the system. Providing the appropriate details on the architecture and implementation of neural networks necessitates some use of jargon, allowing our approach to be compared to other modeling work. We have aimed to limit the use of such jargon to the paragraphs that follow.

The data were randomized and transformed using Standard Scaling, which standardizes measurements by removing the mean and scaling to unit variance using (1):

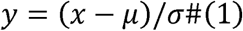

where *μ* is the mean of the training samples; σ is the standard deviation of the training samples. Standard scaling was conducted using the StandardScaler function from scikit-learn [33]. Note that we do not conduct Standard Scaling on the whole dataset at once but in two phases. The scaling model is fitted on the training data. Then, the fitted model is used to transform the training and test sets separately to ensure that there is no information (such as effects on the total mean and standard deviation) from the test set on the training set. This can occur when data are transformed using information from the test and training set combined.

Figure 1 shows the design of the neural network employed in this study. It is a feed-forward neural network (DNN), that conducts multiclass classification. This type of neural network processes data sequentially from the input layer, which is the first layer of the model, to the hidden layers, which are the intermediate layers of the model. It is designed for a multiclass classification as the output classes of the network are three, namely the three PPA variants (an alternative type is the binary classification).

**Figure 1.**
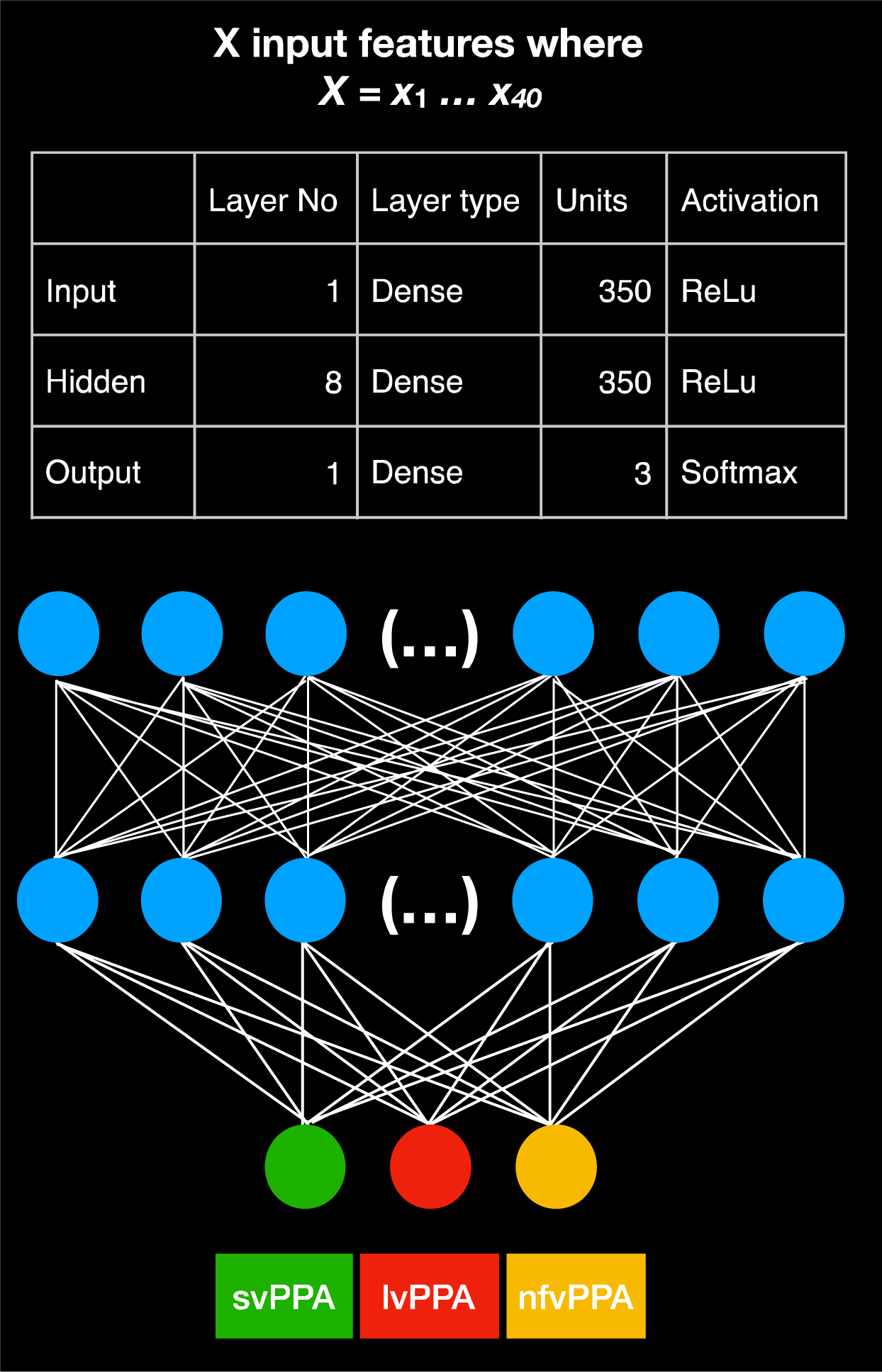
Neural Network Architecture. Structure of the neural network designed for the study and feature properties, including the number of input features employed, the type and number of units and activation functions for the input, hidden, and output layer. The first layer on top is the input layer and consists of 350 units; 8 layers in the middle containing 350 units are hidden layers, and the final layer contains only three units; here with different colors, when the green is activated it corresponds to the svPPA variant, when the red is activated it corresponds to the lvPPA variant, and when the yellow unit is activated it corresponds to the nfvPPA variant.

The prediction of the variant is provided by the last layer a.k.a., output layer. All layers in our model contain units or nodes that are interconnected. The input layer consists of 350 dense units. A Rectified Linear Unit (ReLu) was selected as the activation function of the input and the hidden layers. A ReLu is a mathematical function that returns 0 if it gets any negative input but if it gets a positive value, it will returns that value back, f(x) = max(0, x). ReLu activations have the advantage that compute and converge faster than other activation functions [34]. The output layer does not contain a ReLu activation but a softmax activation function that provides an output, which is either 0 or 1 to facilitate the three-class classification of the PPA variants [35].

We compiled the model using a Root Mean Square Propagation (RMSProp) optimizer [36]. The RMSProp optimizer is a mathematical function that adapts the learning rate for each of the parameters. The learning rate is the step size made towards minimizing the loss function of the network. The RMSProp divides the learning rate for a weight by a running average of the magnitudes of recent gradients for that weight (i.e., mean square). The advantage of RMSProp is that it displays an outstanding adaptation of the learning rate. The loss function we employed was set to categorical cross-entropy, which is most suitable for multiclass classification. A higher value for the loss function implies a greater error for the predictions of the model. We fitted the network batch size set to 32. The model was trained and evaluated eight times, in an 8-fold cross-validation task (see the section on <u>Model comparison and evaluation</u>). In each fold of the crossvalidation, the neural net was trained and evaluated for 30 epochs (an epoch is a single evaluation of the data from the neural network), using a 80% vs. 20% validation split that divides data into an 80% training set and a 20% evaluation set. This evaluation demonstrates the learning progress of the neural network.

### Model comparison and evaluation

To estimate how well the model performs and how it generalizes on unknown data, we employed established evaluation methods that select data from a set of participants that were subtyped by clinicians manually to train the model and a different set of participants to evaluate the model. During the training phase the input of the model are the measurements from the cookie theft picture description task produced by a certain patient. The model is trained multiple times on the measurements to identify the patterns that correspond to the PPA variant of that patient, as was estimated from the clinical neuropsychological assessment, MRI scans, etc. In other words, the goal of the training phase is to find speech and language patterns from the input measurements that characterize the provide PPA variant of the patient. The training phase is followed by the evaluation phase of neural network model. For the evaluation, the input of the model are data from patients that were not employed during the training phase, the PPA variant of these new patients is not provided to the model but it is withheld to evaluate the model predictions. Based on its training and using the new input, the neural network predicts the PPA variant of the new patient. For example, the neural network predicts based on the input data that a patient corresponds to the lvPPA variant; if that patient has been classified during neuropsychological assessment as lvPPA, then we score the assessment as correct; if it is incorrect, then the prediction is evaluated as incorrect. The accuracy of the model is based on the count of correct predictions, true positives, true negatives, and incorrect predictions, false positive and false negatives. The true positive and true negative are the outcomes where the machine learning model correctly predicts the positive and negative class correspondingly. Furthermore, a false positive or a false negative is an outcome where the model predicts the positive and negative classes incorrectly.

To make the best use of our collected data, we employed an *eight-fold* group cross-validation method that allows us to use all the data as training data and all the data as test data but at different 8 different training/evaluation phases, so that we are always employing data from different patients to train the model and data from different patients to evaluate the model. The *eight*-*fold* group cross-validation method splits the randomized data into eight folds and trains and evaluates the machine learning models eight times. During each training session, *seven folds* are employed for training and *one fold*, which contains data from participants that are not in the training folds, is employed as an evaluation set. Therefore, the machine learning models were trained on different folds of data in the training set and evaluated on different test data from unknown participants during evaluation. This evaluation ensures that data are randomized for splitting and that there are always different participants in the training and test sets.

To evaluate the models, we employed the following metrics: *accuracy, precision*, and *recall. Accuracy* is the total sum of correct predictions divided by the total number of both correct and incorrect predictions:

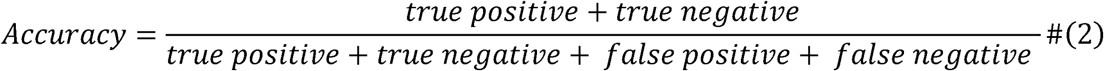

Precision is the result of division of the true positives with the sum of true positives and false positives (see formula 3). Recall (a.k.a., sensitivity) is the result of dividing the true positives with the sum of true positives and true negatives (see formula 4).

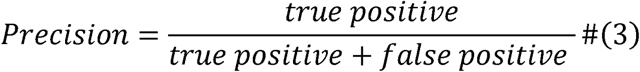

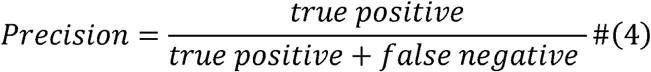

Finally, the *F1 score* is the weighted average of the precision and recall, and ranges between 0 and 1 (see formula 5). The *F*_*1*_ *score* can offer a more balanced estimate of the outcome than the accuracy.

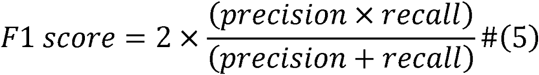

All models were implemented in Keras (Chollet & others, 2015) running on top of TensorFlow (Abadi et al., 2016) in Python 3.6.1.

### Model optimization and hyperparameter tuning

For the selection of the final neural network architecture, we tested several neural network architectures by varying both the number of hidden layers, the number of units per layer, the dropout [37], the activation methods, and the batch size. DT models are provided here as a comparison model and their output is reported without optimizations. We evaluated the SVM models with both linear and non-linear kernels and optimized the models for the number of kernels by running the SVM models with 1 - 300 kernels. The SVM model contains 14 non-linear kernels, which provided the best results in SVM optimization. We evaluated the RF models by optimizing for the number of trees from 1 - 300 trees. The best RF model was the one with 14 trees. Note that the minimum split number was set to two.

